# VTA dopamine neurons are hyperexcitable in 3xTg-AD mice due to casein kinase 2-dependent SK channel dysfunction

**DOI:** 10.1101/2023.11.16.567486

**Authors:** Harris E. Blankenship, Kelsey A. Carter, Nina T. Cassidy, Andrea N. Markiewicz, Michael I. Thellmann, Amanda L. Sharpe, Willard M. Freeman, Michael J. Beckstead

## Abstract

Alzheimer’s disease (AD) patients exhibit neuropsychiatric symptoms that extend beyond classical cognitive deficits, suggesting involvement of subcortical areas. Here, we investigated the role of midbrain dopamine (DA) neurons in AD using the amyloid + tau-driven 3xTg-AD mouse model. We found deficits in reward-based operant learning in AD mice, suggesting possible VTA DA neuron dysregulation. Physiological assessment revealed hyperexcitability and disrupted firing in DA neurons caused by reduced activity of small-conductance calcium-activated potassium (SK) channels. RNA sequencing from contents of single patch-clamped DA neurons (Patch-seq) identified up-regulation of the SK channel modulator casein kinase 2 (CK2). Pharmacological inhibition of CK2 restored SK channel activity and normal firing patterns in 3xTg-AD mice. These findings shed light on a complex interplay between neuropsychiatric symptoms and subcortical circuits in AD, paving the way for novel treatment strategies.

## INTRODUCTION

98% of Alzheimer’s disease (AD) patients experience neuropsychiatric symptoms ranging from anxiety to clinical depression^1^. These conditions, which cannot be fully accounted for by the cortical and hippocampal degeneration typically associated with AD, are consistent with subcortical dysregulation. In humans, AD is strongly co-morbid with Parkinson’s disease (PD)^2^, and extrapyramidal motor symptoms are commonly observed^3^, which could suggest a role for mesencephalic dopamine (DA) in AD pathology. Recent findings from preclinical AD models include fewer tyrosine hydroxylase positive (TH^+^) neurons commensurate with decreased dopamine release^4^, aberrant firing patterns of ventral tegmental area (VTA) DA neurons^5^, and increased dopamine receptor expression, particularly in the VTA^6^. However, investigations into single-neuron activity and changes in cellular physiology during disease progression are sparse and predominantly limited to cortical and hippocampal studies of amyloid-only models during prodromal stages of the disease.

VTA dopaminergic neuron projections to the cortex contribute to episodic memory formation^7–9^ while subcortical projections, particularly to the nucleus accumbens (NAc), encode reward, motivation, and aversion^10,11^. While cognitive tests^4^ and locomotion^6^ are common in the preclinical AD literature, reward learning is largely unexplored. Reward-based behaviors rely on both tonic extracellular DA regulated by basal firing of dopaminergic neurons and phasic release of DA mediated by burst firing^12–14^. In the substantia nigra pars compacta (SNc), autonomous firing of DA neurons is central to the initiation of degeneration in PD^15^.

Conversely, in AD, the mesocorticolimbic system (i.e., the VTA) seems to be preferentially affected over the nigrostriatal pathway^4,16,17^. Distinct biophysical mechanisms underlie firing patterns in the SNc and VTA^13^, potentially contributing to the selectivity of neurodegenerative diseases within DA neuron subpopulations. Ion channels govern the electrophysiological properties of excitable cells, and altered channel function and action potential generation have been observed in the hippocampus of AD models^18,19^. As single neurons are the computational units of the brain, determining physiological alterations of individual cell populations in early AD is central for understanding disease pathology and for identifying potential therapeutic targets.

Here, using the amyloid + tau-based triple-transgenic 3xTg-AD mouse model (3xTg)^20,21^, we describe deficits in reward-based learning that prompted an investigation into firing and ion channel function in single VTA DA neurons. We observed hyperexcitability of DA neurons caused by decreased small-conductance calcium-activated potassium (SK) channel currents in 3xTg compared to age-matched control mice. This was driven by SK-bound casein kinase 2 (CK2), which decreases the calcium binding affinity of calmodulin (CaM) and effectively reduces SK channel activation by calcium. Importantly, pharmacological inhibition of CK2 was sufficient to restore basal firing rate and regularity in 3xTg DA neurons. These results reveal a noncanonical deficit in dopaminergic behavior and single cell physiology that hinges on hyperactivity of a single AD-associated enzyme, CK2, implicating it as a potential therapeutic target for AD.

## RESULTS

### Reinforcement learning is impaired in 3xTg-AD mice

3xTg mice carry homozygous transgenes for human APP_Swe_, tau_P301L_, and PS1_M146V_, all of which are linked to familial AD. These mice accumulate both extracellular beta-amyloid (Aβ) and intracellular hyperphosphorylated tau^20^, enabling exploration into neuropsychological abnormalities and the underlying physiological mechanisms. AD patients suffer from deficits in motivation-based appetitive behaviors that could indicate a central role for VTA DA neurons^16^. To assess whether 3xTg mice display behavioral impairment in operant reward learning, we assessed acquisition of an operant task to gain access to palatable pellets in non-food restricted 6- and 12-month-old 3xTg and wildtype (WT) mice (Figure 1a), a task previously shown to rely on NAc DA^22^. All WT mice tested at both ages successfully acquired the operant task. However, 3xTg mice at both ages exhibited slower learning of the task, and several mice were unable to learn the task at all, especially at the 12-month time point (Figure 1b). Raster plots of nose poke responding on day 1 of training for individual mice that eventually learned the task show less responding over the 3-hour session compared to WT mice, although some 3xTg mice did exhibit responding similar to WT mice (Figure 1c). Among the mice who acquired the task, there was no main effect of age or genotype on the mean number of pellets earned during a session (Figure 1d), or on motivation to gain access to the reinforcer, as measured by progressive ratio breakpoint (Figure 1e). Due to these clear deficits in reward learning in 3xTg mice, we aimed to probe the physiological properties of single DA-releasing neurons in the VTA.

**Figure 1.**
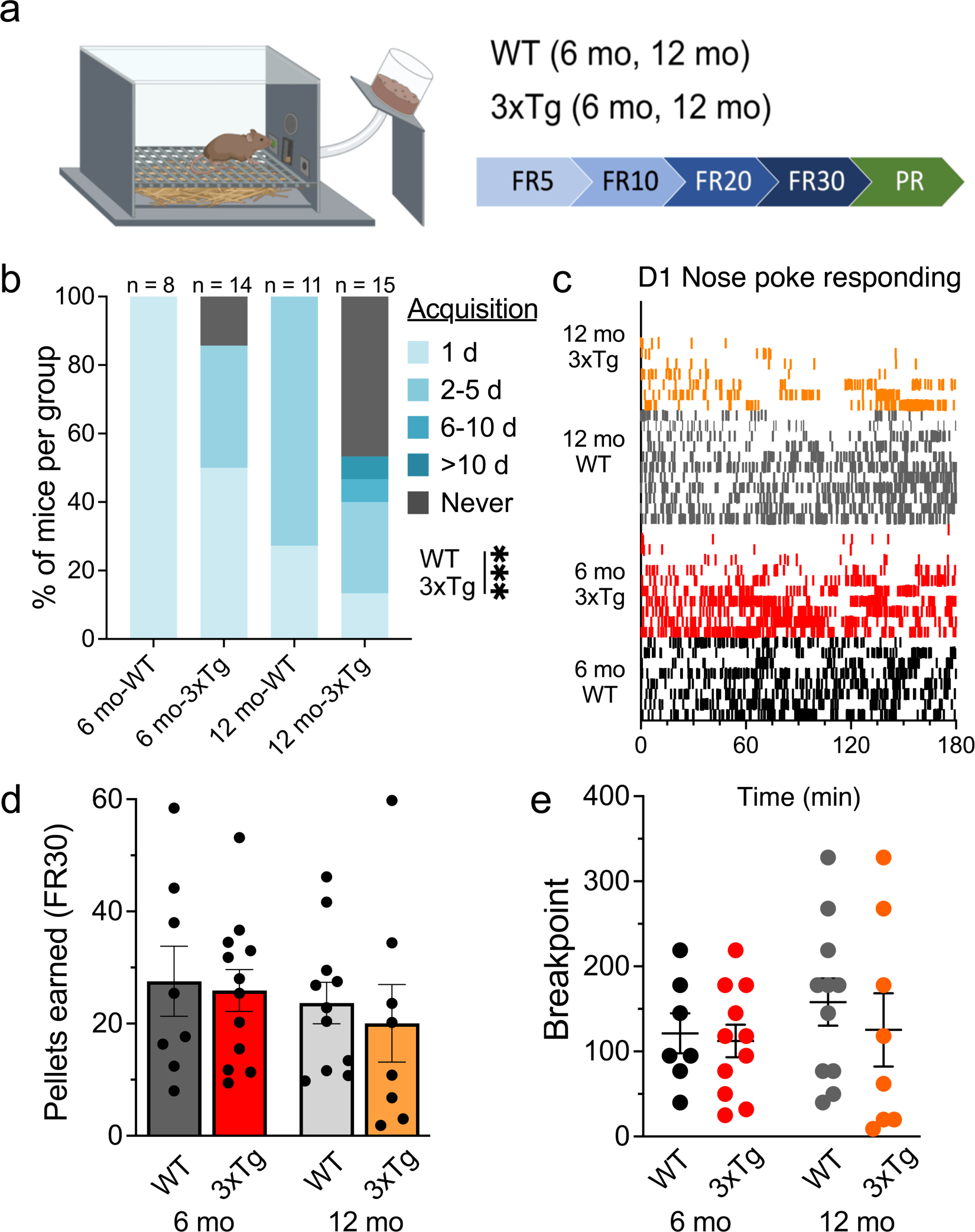
Instrumental reward learning is age-dependently impaired in 3xTg mice. **a.** Schematic of procedure used for operant self-administration of palatable food pellets. **b.** 3xTg mice acquired operant self-administration of palatable pellets more slowly than WT mice (main effect, *P* = 0.0007, F_1,44_ = 13.27), and older mice learned the task more slowly than younger mice (main effect, *P* = 0.0156, F_1,44_ = 6.34). **c.** Raster plots showing correct side nose poke responding (vertical (cks along the x-axis) on day 1 of training for individual mice (by row) that eventually acquired self-administration. While there is considerable variation within the 3xTg groups, overall, they show less responding on day 1 compared to WT mice. **d.** Although there was a significant effect on learning the operant task, the number of pellets earned at FR30 did not significantly differ by age (*P* = 0.337) or genotype (*P* = 0.600). **e.** The breakpoint during the PR test session was not different by age (*P* = 0.402) or genotype (*P* = 0.483).

### 3xTg DA neurons exhibit increased firing rate and irregularity

While dopamine release can occur independent of somatic activity^23,24^, alterations in intrinsic DA neuron physiology are sufficient to affect reinforcement learning^25^. To examine the physiological properties of VTA DA neurons across the lifespan and in response to amyloid and tau expression, patch clamp recordings were made in brain slices from WT and 3xTg mice at 3, 6, 12, and 18 months of age. In rodent brain slices, VTA DA neurons spontaneously fire in a slow (0.5-4 Hz), pacemaker-like pattern^26–28^. Using the cell attached configuration (Figure 2a) to assess spontaneous firing of DA neurons without disrupting intracellular contents, we observed increased spontaneous firing frequency in 3xTg mice compared to age-matched controls (Figure 2b). 3xTg mice also displayed lower firing rhythmicity as indicated by an elevated coefficient of variation (CV) of the inter-spike intervals (ISI; Figure 2c). While the CV of the ISI was high throughout the lifespan, the increase in firing rate exhibited an inverted-U shape, peaking at 12 months before declining to WT frequency at 18 months (Figure 2b). Since the ionic mechanisms that control pacemaking tightly couple to those governing a shift to burst firing VTA DA neurons^12,29^, we next sought to determine how 3xTg neurons responded to electrical input.

**Figure 2.**
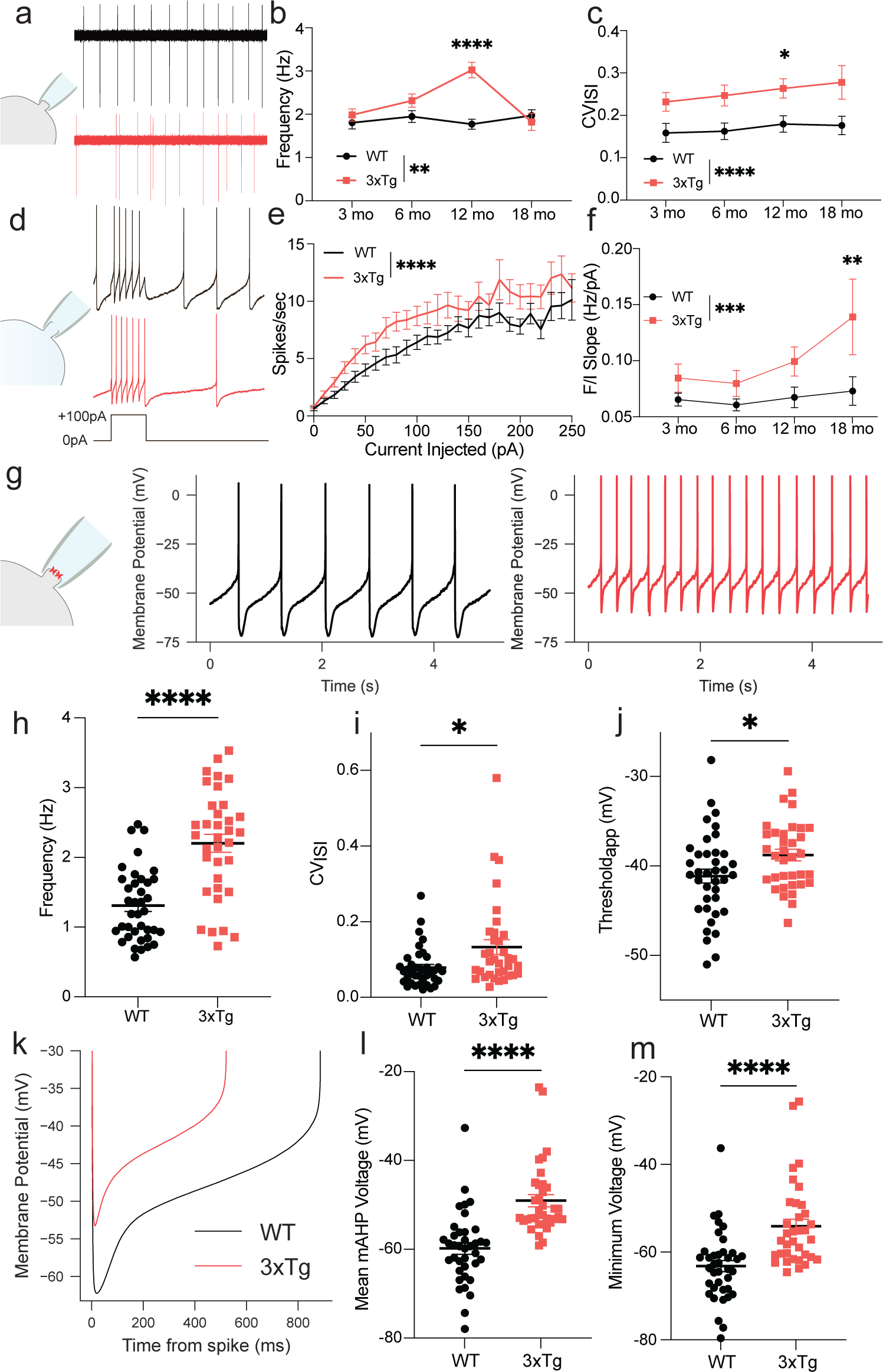
VTA DA neurons from 3xTg mice display increased firing rates and decreased regularity. **a**. Schematic of a cell-aSached recording and 10 s representative traces of spontaneous firing activity from 12 mo WT (black) and 3xTg (red) mice. **b.** Mean spontaneous firing rate for 3, 6, 12, and 18 mo WT and 3xTg DA neurons (N = 576 cells, two-way ANOVA F_genotype_ = 9.236, P=0.0025; F_age_ = 3.878, P=0.0092; F_interaction_ = 5.943, P=0.0005), with the most notable increase in rate at 12 mo (P<0.0001). **c.** Mean coefficient of variation of the ISI (CV_ISI_) collected for the same cells as in **b** (F_genotype_ = 19.64, P <0.0001), and showed significant difference between 3xTg and WT at 12 mo (P=0.0214). **d.** Schematic of a whole-cell recording and representative traces of +100 pA current injection for 1 s in WT (black) and 3xTg (red) mice. **e.** Averaged frequency/current (F/I) curves from 12 mo WT and 3xTg mice (n=24 WT and n=31 3xTg, F_genotype_ = 54.56; P<0.0001, F_current injection_ = 18.48, P<0.0001). **f**. Hypersensitivity of 3xTg DA neurons measured as the slope of the F/I curve (0-60 pA, N = 222 cells, F_genotype_ = 13.71, P=0.0003; F_age_ = 2.692, P=0.0471), with a significant increase in sensitivity at 18 mo in 3xTg DA neurons (P=0.0044). **g**. Schematic of perforated-patch gramicidin recording and representative traces from spontaneously firing DA neurons from WT (black) and 3xTg (red) mice. **h**. Perforated-patch recordings show increased spontaneous firing frequency in 3xTg neurons (n = 38 WT and 35 3xTg neurons, two-tailed t-test, P<0.0001), and **i**. increased CV_ISI_ (two-tailed t-test, P=0.0363). **j**. Depolarized apparent threshold in 3xTg DA neurons (two-tailed t-test, P=0.0247). **k**. Grand averages of ISI voltage trajectories for WT and 3xTg cells. **l**. Depolarized mean medium aeer-hyperpolarization voltage (mAHP, two-tailed t-test, P<0.0001), and **m**. minimum ISI voltage (two-tailed t-test, P<0.0001) in 3xTg DA neurons.

*In vivo*, glutamatergic afferents drive VTA DA neuron burst firing in response to reward-predictive cues^30^. These synaptic connections are severed in *ex vivo* brain slice preparations, and DA neurons do not typically burst under basal conditions. Somatic current injection can mimic burst activity and provides insight into how dopamine neurons respond to excitatory input^12^. Using whole-cell current-clamp recordings, we next applied a series of current steps (0-250 pA) to generate frequency-current (F-I) curves for WT and 3xTg mice of all four ages (Figure 2d,e). 3xTg dopamine neurons were hypersensitive to current injection, as quantified by the slope of F-I curves, or gain (Figure 2f). To further assess the effects on firing, we employed the minimally-invasive gramicidin perforated patch recording technique to produce stable, long-duration recordings while maintaining electrical access to the cell. Focusing on 12-month-old mice, we again observed a robust increase in frequency and irregularity in 3xTg DA neurons (Figure 2g,h,i). Additionally, in this recording configuration, we were able to detect a significant depolarization of the apparent spike threshold in 3xTg DA neurons (Figure 2j), but no change in the spike width (data not shown, *P*=0.9781), suggesting high threshold calcium channels and action potential repolarization mechanisms are unaltered^31^.

To test whether altered firing generalized to other dopaminergic neurons, we also recorded from an additional located in the SNc. This neuron type exhibits spontaneous firing activity that is governed by similar ion channel conductances to those of VTA neurons^32^. In the SNc we detected no alterations in spontaneous firing rate or CV of ISI in 12-month-old 3xTg mice (Extended Figure 1b,c,d). Furthermore, evoked firing (Extended Figure 1e,f,g) was similar to that of WT mice. These findings are consistent with a prior report of preserved SNc DA neuron function in an amyloid-only model of AD^4^. Thus, VTA dopamine neuron firing appears to be selectively disrupted among DA neurons in 12-month-old 3xTg mice.

The biophysical properties responsible for slow spontaneous firing of midbrain DA neurons are extremely well characterized. In the VTA, the two major outward currents are carried by A-type potassium (Kv4.3)^34–36^ and SK (largely SK3) channels^29,37–39^, which interact across the ISI to maintain a stable firing cycle. We sought to determine how ISI dynamics in VTA DA neurons differed between WT and 3xTg mice. Inter-spike voltage trajectories were collected and averaged for both genotypes (Figure 2k). The mean medium afterhyperpolarization (mAHP) voltage was depolarized in DA neurons from 3xTg mice (Figure 2l) as was the minimum voltage of the ISI (Figure 2m). The mAHP is governed largely by action potential mediated influx of calcium and subsequent activation of SK channels^40^. Indeed, the reduced mAHPs and minimum ISI voltages in 3xTg mice were highly reminiscent of data we recently described in the SNc following a partial block of SK channels using apamin^41^, and a depolarized threshold may be secondarily linked to SK channels through sodium channel inactivation^38,42^. Thus, we aimed to determine whether SK channel activity is altered in 3xTg VTA DA neurons.

### SK channel dysfunction underlies aberrant firing

SK channels are activated by calcium influx through voltage-gated calcium channels and subsequent calcium-activated calcium release^43,44^. SK activity can be measured as an afterhyperpolarization (tail) current in SNc DA neurons following a depolarizing voltage step^42,45^, but this has not been previously reported in adult VTA. We therefore applied a saturating concentration of the selective SK channel inhibitor apamin (100 nM)^46^ to assess its effect on the tail currents in VTA DA neurons (Extended Figure 2). Apamin rapidly and consistently decreased the amplitude of the tail current from an average of 329.4 to 109.8 pA (Extended Figure 2b,c). We therefore assessed tail currents using the same protocol (100 ms depolarizing step from −72 to −17 mV) in DA neurons from 3-, 6-, 12-, and 18-month-old WT and 3xTg mice (Figure 3a). We observed markedly smaller tail current amplitudes (Figure 3b) and area under the curve (Figure 3c) in 3xTg mice. Consistent with earlier data, there was a significant age effect on both measures (Figure 3b-c). Notably, we did not detect any change in the tail current AUC or maximal amplitude in SNc DA neurons (Extended Figure 1h,i), again demonstrating selectivity for VTA among DA neurons.

**Figure 3.**
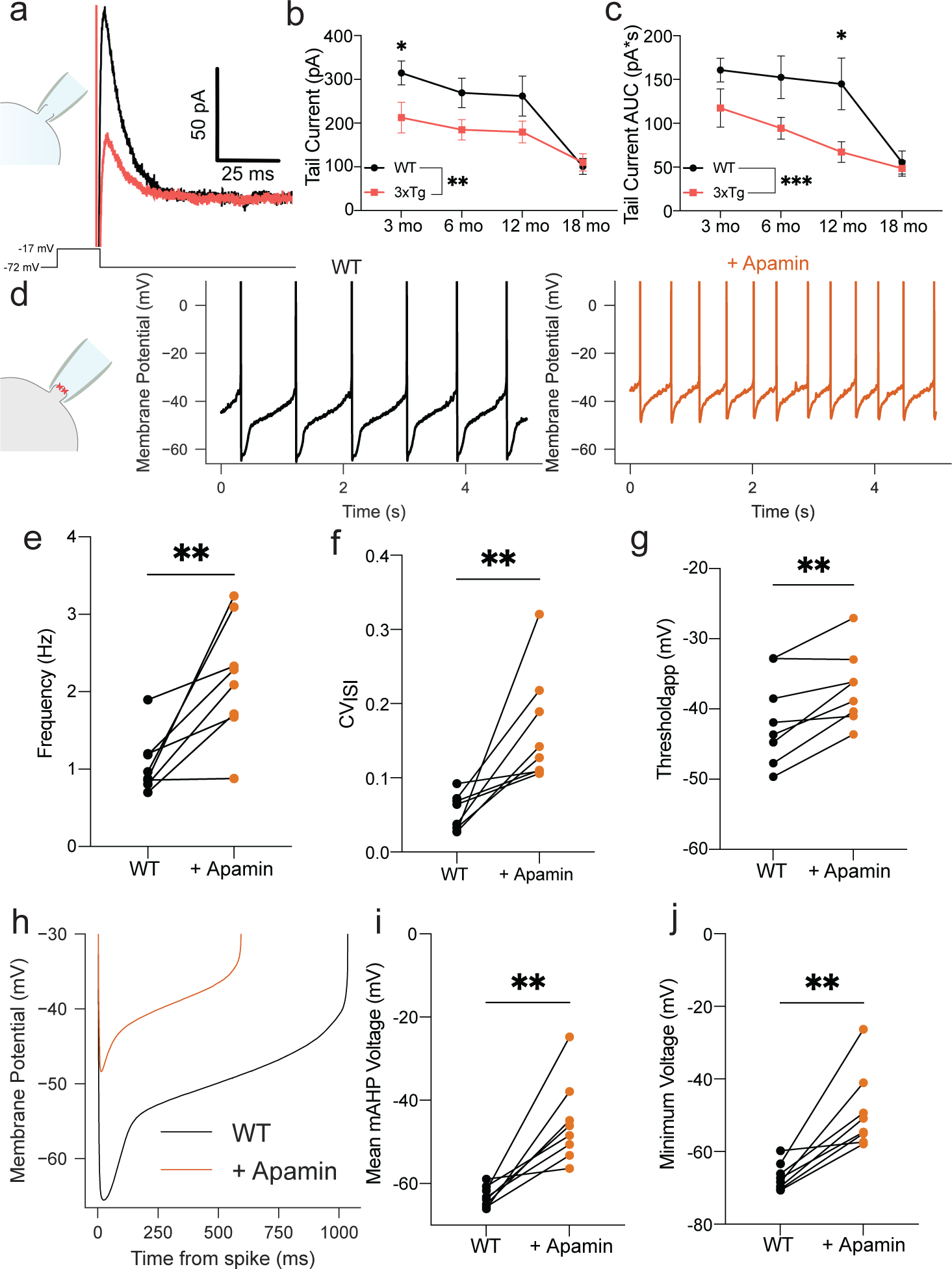
SK channel inhibiGon in WT neurons recapitulates the 3xTg phenotype. **a.** Representative traces of the tail current in response to a 100 ms depolarization step to −17 mV from V_hold_ (−72 mV) in the whole-cell configuration. **b.** Reduced tail current maximal amplitude (N = 251 cells, two-way F_genotype_ = 8.743, P=0.0034; F_age_ = 10.57, P<0.0001) and **c.** area under the curve (AUC, F_genotype_ = 13.79, P=0.0002, F_age_ = 10.03, P<0.0001) in 3xTg DA neurons. **d**. Perforated-patch recordings from spontaneously firing WT cells in control ACSF (black) and in the presence of apamin (1-3 nM) (orange). **e**. Increased firing frequency (paired two-tailed t-test, P=0.0067) and **f.** CV_ISI_ (paired two-tailed t-test, P=0.0088) in presence of apamin. **g.** Depolarized apparent spike threshold in response to apamin (paired two-tailed t-test, P=0.0050). **h.** ISI voltage trajectory averages. The effect of apamin is reminiscent of the trajectories from 3xTg neurons (panel **1k**). **i**. Depolarized mAHP (paired two-tailed t-test, P=0.0022) and absolute minimum ISI voltages (paired two-tailed t-test, P=0.0018) in apamin.

To determine whether partial block of SK channels recapitulates the physiological abnormalities seen in 3xTg mice, we applied a low concentration of apamin (1-3 nM) to spontaneously firing VTA DA neurons from 12-month-old WT mice in the perforated patch configuration (Figure 3d). We observed an increase in firing rate (Figure 3e) and CV of ISI (Figure 3f) similar to what is observed in 3xTg neurons. Moreover, the ISI dynamics of spontaneously firing VTA DA neurons mimicked those of 3xTg VTA DA neurons (Figure 3h), with mean mAHP voltage (Figure 3i), minimum ISI voltage (Figure 3j), and spike threshold (Figure 3g) all depolarized. These data are consistent with SK channel involvement in the physiological abnormalities of VTA DA neurons in 3xTg mice.

We also determined whether alterations extended to the other major rate-limiting potassium current, mediated by A-type potassium channels (I_A_). I_A_ is critical for the slow depolarizing ramp across the ISI due to its inactivation voltage and slow decay time constant^34,35,47^. We used whole-cell voltage-clamp recordings to isolate I_A_ using a series of voltage steps (Extended Figure 3a)^35,36,47^. Interestingly, neither I_A_ amplitude (Extended Figure 3b) nor decay time constant (Extended Figure 3c) differed between WT and 3xTg VTA DA neurons, nor was there a difference in the SNc (Extended Figure 1j,k), suggesting I_A_ is unaltered in midbrain DA neurons of 3xTg mice.

### Enhancing SK function normalizes firing of 3xTg DA neurons

Positive allosteric modulators of SK channels have previously been shown to safeguard neurons from neurotoxic insults in degenerative models^48,49^. We next tested the sensitivity of VTA DA neurons by applying the selective SK channel positive allosteric modulator NS309 (1 µM) in perforated-patch recordings of WT (Figure 4a) and 3xTg VTA DA neurons (Figure 4b)^41,50^. NS309 reduced firing rate in WT, but not 3xTg neurons (Figure 4c). However, it had no significant impact on the CV of the ISI in either genotype (Figure 4d), contrary to our recent findings in SNc neurons^41^. When we further scrutinized ISI dynamics, we observed that WT neurons (Figure 4e) were more sensitive than 3xTg (Figure 4f) neurons; NS309 hyperpolarized the average mAHP (Figure 4g) and the minimum ISI voltage (Figure 4h) in WT but not 3xTg VTA DA neurons. These data suggest that SK channels in 3xTg mice might be less sensitive to modulation by NS309.

**Figure 4.**
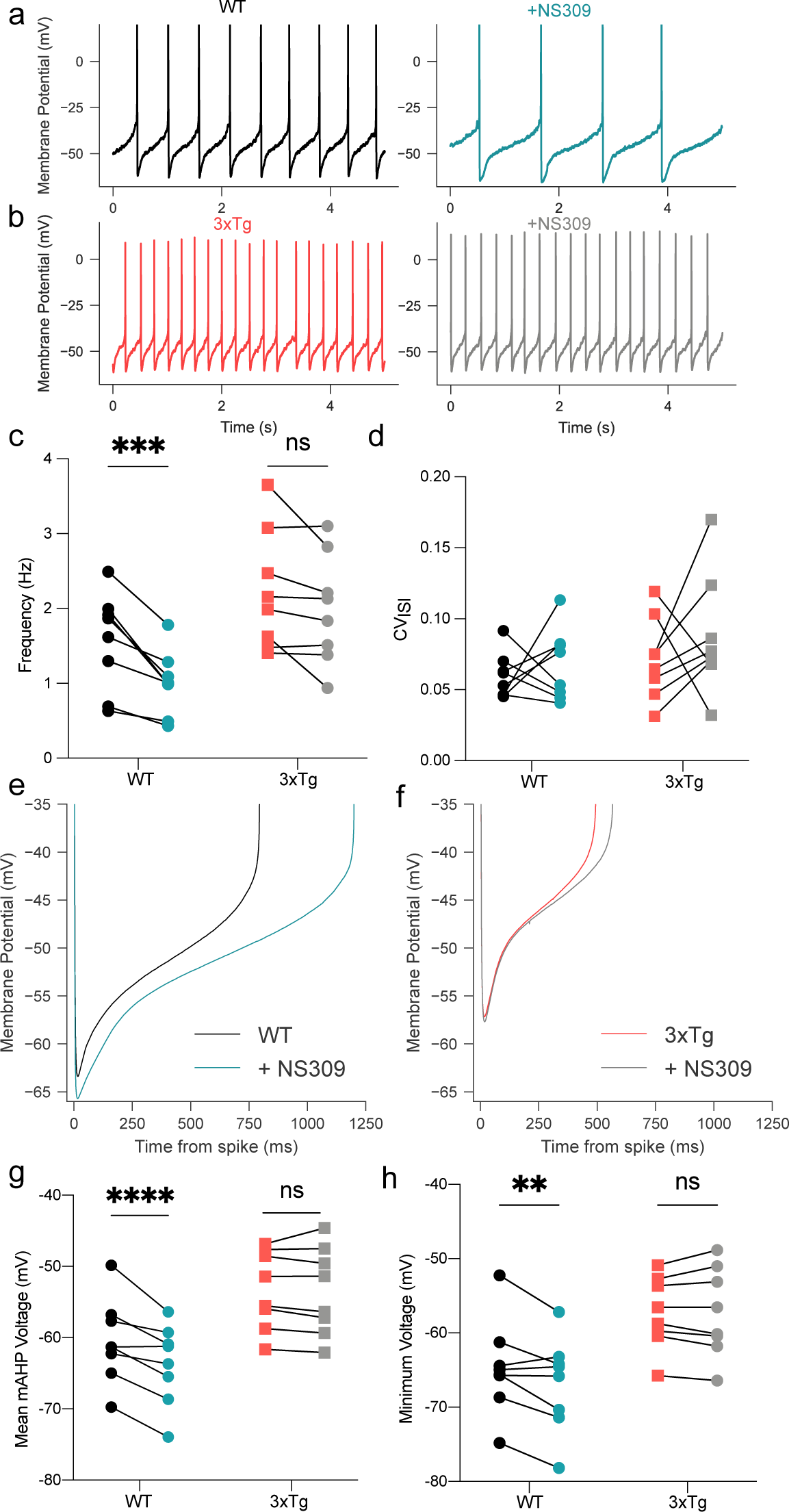
NS309 efficacy is decreased in 3xTg DA neurons. **a**. Representative traces of WT DA neuron pacemaking in control ACSF (black) and in the presence of NS309 (1 µM) (blue). **b**. Representative traces of 3xTg DA neurons in control ACSF (red) and in NS309 (gray). **c**. Firing frequency was decreased by NS309 (two-way RM ANOVA, F_NS309_ = 15.27, P = 0.0058; F_genotype_ = 5.872, P=0.0459; F_NS309 x genotype_ = 6.604, P=0.0370) in WT (P=0.0007) but not significantly in 3xTg (P=0.0554) neurons. **d**. NS309 did not affect CV_ISI_. **e**. Averaged ISI voltage trajectories in WT neurons before (black) and aeer NS309 (blue) and **f**. in the 3xTg group before (red) and in the presence of NS309 (gray). **g**. NS309 hyperpolarizes mean mAHP voltages in WT (P<0.0001), but not 3xTg (P=0.7795) neurons (two-way RM ANOVA, F_NS309_ = 9.963, P=0.0160, F_genotype_ = 14.89, P=0.0062, F_NS309 x genotype_ = 43.200, P=0.0003). **h**. NS309 hyperpolarizes minimum ISI voltages (F_genotype_ = 11.06, P=0.0127; F_NS309 x genotype_ = 11.94, P=0.0106) in WT (P=0.0037) but not 3xTg (P=0.9988) neurons.

SK channels function as multi-protein complexes that continuously associate with CaM as a calcium sensor^51^. Ca^2+^ binding to CaM produces a conformational change in the channel that results in pore opening and potassium efflux^52^. NS309 augments the apparent calcium sensitivity of SK channels by binding to the channel-CaM interface, allosterically enhancing phosphatidylinositol 4,5-bisphosphate (PIP2) recruitment to promote channel opening^53,54^. SK channels can be dynamically phosphorylated at threonine 79^55^, and upon phosphorylation the NS309 binding affinity is decreased^54^. Our findings point toward the possibility that an additional subcellular participant might phosphorylate CaM to alter basal and modulated SK channel activity in 3xTg DA neurons. To investigate the expression of SK channel interactors in DA neurons from WT and 3xTg mice, we turned to a molecular screening technique.

### Molecular interactors of SK channels

To identify endogenous SK channel interactors, we coupled patch clamp electrophysiology with single-cell transcriptomics (Patch-seq)^56^. While the technique was originally developed to profile neuronal subpopulations^56,57^, we posited it could be used as a screening assay akin to single-cell or single-nucleus RNA sequencing. The latter techniques have proven instrumental in detecting genes responsible for vulnerability in both post-mortem brain tissue of human patients and animal models of disease^58,59^. Patch-seq has been used previously in this context to study pancreatic β-cells in a model of type-2 diabetes^60^, in an *in vitro* model of schizophrenia^61^, and in a model of corticospinal tract degeneration^62^, all of which returned potential mechanistic targets of physiological dysfunction, but this technique has not been previously used in brain sections from a preclinical disease model (including those for Alzheimer’s or other neurodegenerative diseases).

Using slightly modified procedures to minimize artifacts and cell swelling^63^, we obtained patch clamp recordings in 12-month-old WT and 3xTg mice (Figure 5a). We again observed reduced SK currents in 3xTg mice and extracted a total of 48 cells from the two groups. 4,824 genes passed criteria for expression and transcriptome profiles, and WT and 3xTg generally separated in the 1^st^ component of PCA space (70% of explained variance) (Figure 5b). The grouping also indicated a larger heterogeneity in 3xTg neurons, both by transcriptomic profile (PCA) and by physiological measure (SK current). 34 cells generated libraries sufficient for sequencing and passed QC. Using thresholds FDR < 0.01 and a |fold change| > 1.25, there were 3387 differentially expressed genes (DEGs), most upregulated (3373 higher in 3xTg than in WT, and 14 lower in 3xTg than in WT) (Figure 5C). DEGs were then assessed for correlation with individual cell tail current AUC values (|Euclidian| 0.6-1.0). All 14 downregulated DEGs were positively correlated with tail current AUC, while upregulated DEGs demonstrated 111 positively correlated with tail current AUC and 31 negatively correlated (Figure 5D). DEGs were also subjected to molecular function gene ontology (GO) analysis. These analyses indicated increased ion channel binding (GO:0044325) driven by genes including *Calm1*, which codes for the SK channel calcium sensor calmodulin (Figure 5E). From the DEGs, example transcripts of interest included *Grk4*, which encodes the G protein-coupled receptor kinase 4 (positively correlated with tail current AUC) and a number of channels and receptors such as *Cacnd1a2*, which codes for the α_2_δ-1 subunit of voltage gated calcium channels. Indeed, there was not a significant change in the Kcnn3, the gene coding for SK3 (*P*=0.0553) suggesting channel expression levels do not explain tail current measurements. However, the gene that provided the greatest clue into SK channel dysregulation was *Csnk2a1*, a gene coding for casein kinase 2 (CK2), whose expression was also negatively correlated with tail current AUC. CK2 is a ubiquitous kinase and has been a focus of the cancer metastasis field for decades^64^. However, CK2 also has a critical central nervous system role as a kinase that binds to SK channels and regulates their calcium sensitivity^51,55^. We thus sought to determine its role in SK channel regulation in 3xTg mice.

**Figure 5.**
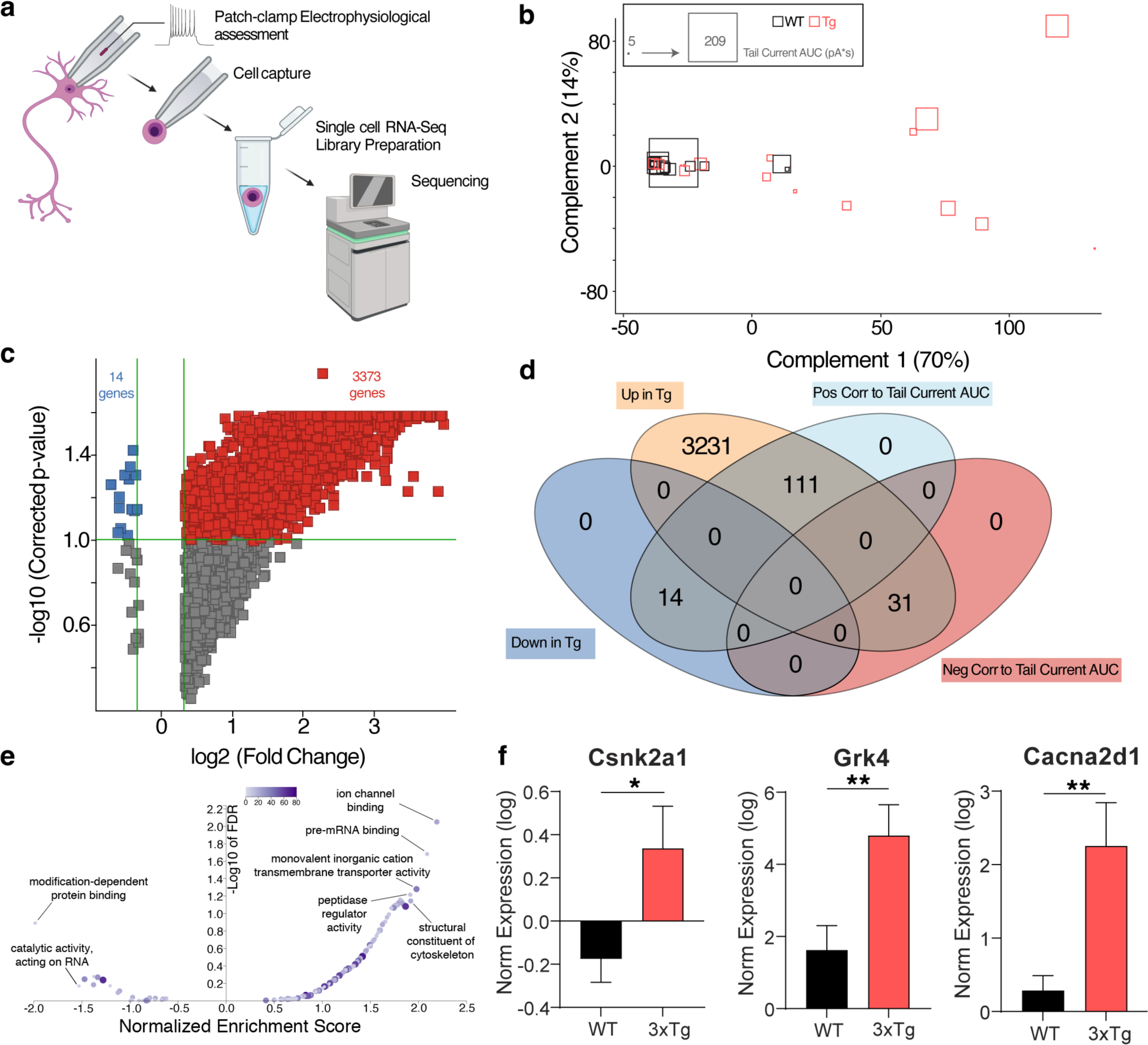
– Patch-seq analysis indicates up-regulated Csnk2a1 transcripGon. **a.** For Patch-seq, recording from an individual neuron is followed by cell extraction with the recording pipeSe. Cell contents are captured, and each individual cell RNA-Seq library is prepared and sequenced. **b.** Individual cells projected in transcriptome PCA space and sized by Tail Current AUC (pA s) demonstrated some separation of 3xTg and WT. **c.** Volcano plot of differentially expressed genes (t-test, B-H MTC, FDR<0.1, |FC|>1.25). **d.** Venn diagram describing overlap in differentially expressed genes between groups and those correlating to Tail Current AUC. **e.** Volcano plot describing molecular function gene ontology analysis (weighted set cover redundancy reduction) on DEGs indicating increased ion channel binding (GO:0044325) pathways (FDR=0.0089544, Normalized Enrichment Score=2.1932). **f.** Example differentially expressed genes (Csnk2a1, Grk4, and Cacna2d1). *FDR<0.1, **FDR<0.05.

### CK2 is central to DA neuron dysfunction in 3xTg mice

CK2 modulates SK channels in an activity-dependent manner^55,65,66^. Previous work has highlighted SK-associated CK2 activity in a model of *status epilepticus*^67^, another form of hyperactivity. CK2 is upregulated in AD patient brain tissue^68,69^ and animal models of the disease^70^. CK2 hyperactivity has also been previously associated with cognitive decline in 3xTg mice^71^. Further, Aβ stimulates CK2 activity *in vitro*^71,72^ providing a potential mechanistic link between AD pathology and dopaminergic dysfunction.

To test the effects of CK2 phosphorylation on SK channel currents, we incubated brain slices from 12-month-old WT and 3xTg mice for 1-3 hours with 1 µM SGC-CK-2 (SGC, a membrane-permeant CK2-specific antagonist^73^), conducted perforated-patch recordings, and compared them to naive cells (previously presented in Figure 2). SGC significantly reduced the spontaneous firing rate in 3xTg but not WT VTA DA neurons (Figure 6a,b,c). The CV of ISI was also substantially reduced only in 3xTg VTA DA neurons, and not in WT cells (Figure 6d). ISI dynamics were similarly affected (Figure 6e); SGC had no apparent effect on the mAHP in WT cells but hyperpolarized 3xTg VTA DA neurons to near WT levels (Figure 6f). Similar effects were seen for the minimum ISI voltage (data not shown, *P*<0.0001). Additionally, apparent spike threshold was significantly hyperpolarized in SGC-treated 3xTg neurons (*P*=0.0176) suggesting that secondary physiological adaptions can be restored. To confirm the effect of SGC on the tail current, we compared tail current AUC in the presence of SGC to data presented in Figure 3c (12-month timepoint). Consistent with our spontaneous firing data, we detected no effect of SGC on WT neurons, but a significant increase in tail current charge of 3xTg neurons (Figure 6g).

**Figure 6.**
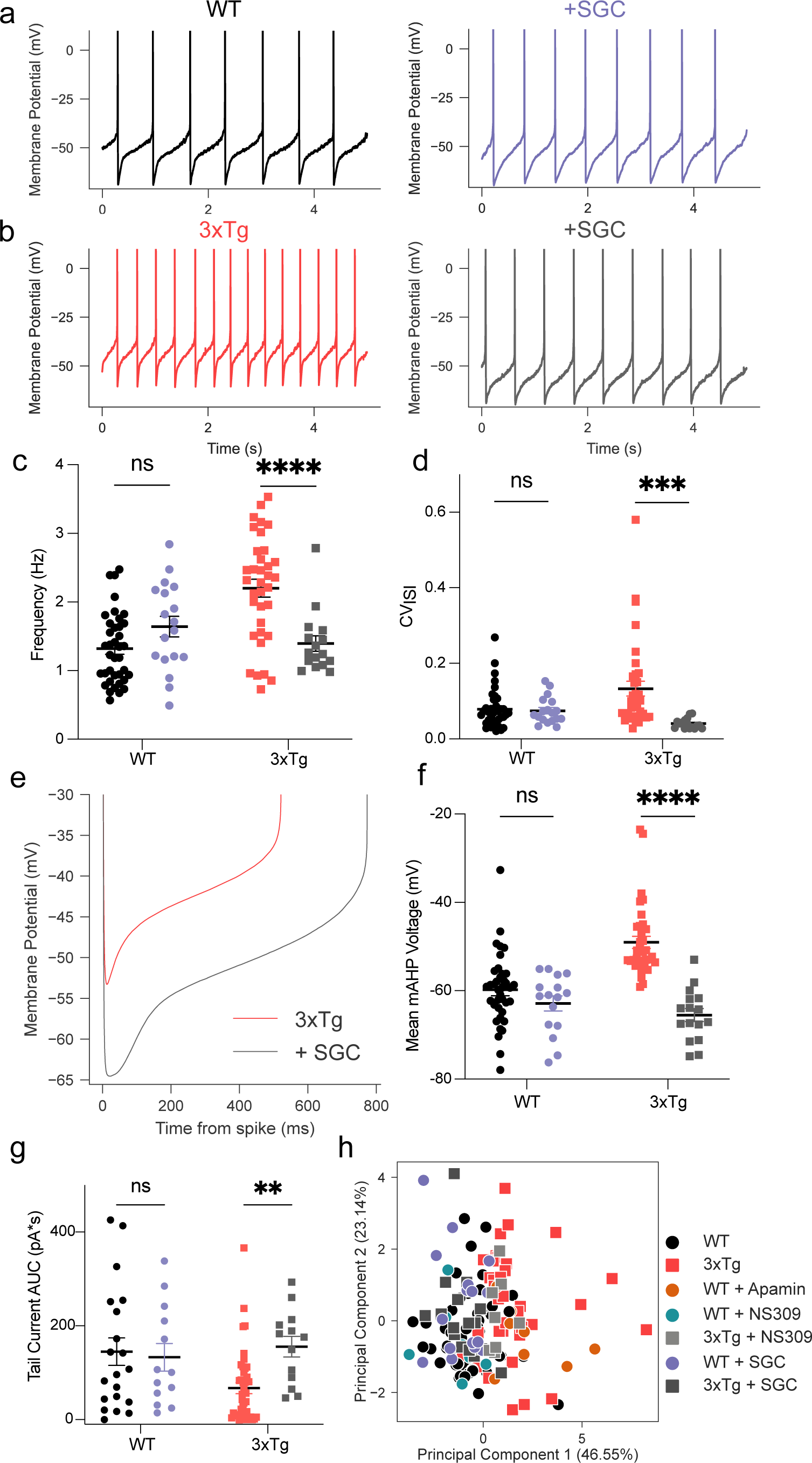
Casein kinase 2 inhibiGon restores firing properGes in 3xTg DA neurons. **a**. Representative firing trace from a naive WT neuron (black) and one pre-incubated with SGC (1 µM, purple). **b**. Representative trace from a naive 3xTg neuron (red) and one pre-incubated with SGC (gray). **c**. SGC decreased firing frequency in 3xTg (P<0.0001) but not WT (P=0.1140) neurons (N = 103 cells; F_genotype_ = 5.988, P=0.0161; F_SGC_ = 3.514, P=0.0637; F_genotype x SGC_ = 18.93, P<0.0001). **d**. SGC decreased CV_ISI_ in 3xTg (P=0.0002) but not WT (P=0.9796) DA neurons (N = 103 cells, F_SGC_ = 9.502, P=0.0026; F_genotype x SGC_= 8.034, P=0.0055). **e**. Averaged ISI voltage trajectories of naïve 3xTg neurons and ones pre-incubated with SGC. **f**. SGC hyperpolarized the mAHP in 3xTg (P<0.0001) but not WT (P=0.3225) neurons (F_genotype_ = 6.349, P=0.0133; F_SGC_ = 36.74, P<0.0001; F_genotype x SGC_ = 17.22, P<0.0001). **g.** Pre-incubation with SGC restored tail current charge in 3xTg (P=0.0098) neurons but had no effect in WT (P=0.9257) neurons (N=84 cells, F_genotype x SGC_ = 4.765, P=0.0318). **h**. Principal component analysis of electrophysiological properties in all groups. The first component describes 46.6% of the variability, and the second describes 23.1% of the variability. 3xTg and WT + apamin cells largely fall to the right, while WT neurons and manipulations as well as 3xTg + SGC segregate to the lee.

Finally, to provide a more global view of electrophysiological properties across experimental groups on which we obtained seven different measures using perforated perforated patch recordings, we used principal component analysis (PCA) for dimensionality reduction to capture variance in a two-dimensional projection. Despite incomplete grouping caused by the heterogeneity of neurons, principal component 1 captured a separation between WT and 3xTg (Figure 6h). Importantly, SGC-treated 3xTg neurons show excellent overlap with WT space. Taken together, these data suggest that CaM bound to plasmalemmal SK channels is hyperphosphorylated in 3xTg VTA DA neurons, resulting in decreased SK channel conductance, increased spontaneous firing rate and irregularity, and hypersensitivity to somatic current injection.

## DISCUSSION

Anecdotal evidence from AD patients suggests a possible role for dopaminergic dysfunction^74,75^, yet no molecular connection between DA neuron pathology and associated neuropsychiatric symptoms has been described. To address this, we first examined mice for their ability to learn operant self-administration of a palatable food reward to test deficits in instrumental learning and motivation. 3xTg mice demonstrated an age-dependent impairment in learning of the task, with no significant change in consumption or motivation in the mice that did learn. We next probed single DA neurons of the VTA and found that a reduction in SK channel conductance produced irregular firing accompanied by an increase in firing rate in slices from 3xTg mice. Finally, we used a combination of whole-cell patch-clamp recordings, RNA sequencing, and pharmacology to implicate CK2 hyperactivity as the likely culprit for the abnormalities.

### Impaired dopaminergic function in 3xTg mice

Converging evidence has recently implicated dopaminergic dysfunction in the neuropsychiatric aspects of AD. Findings using amyloid-only rodent models of AD indicate a DA-dependent decrease in memory formation and reward processing, concomitant with a decrease in TH^+^ cell bodies in the VTA and dopamine release in the NAc^4^. Additionally, hemizygous 3xTg mice exhibit a DA-dependent increase in basal locomotion in post-but not pre-pathological stages^6^. However, although reward-motivated behavior is known to decline in dementia patients^76^, it has not previously been investigated in AD mouse models. Here, we show that non-food-restricted 3xTg mice age-dependently fail to learn an instrumental conditioning task for a palatable food pellet reward. Among the mice that did learn the instrumental task, there was no difference in pellets earned per session or in motivation to gain access to the reinforcer, measured by breakpoint within a progressive ratio test. This may suggest a reduction in the reinforcing properties of the pellets in more affected 3xTg mice, although motivation for the reinforcer was unaffected in the mice that eventually (albeit more slowly) learned the task. These data are congruent with clinical data suggesting heterogeneity in disease progression and penetrance, as well as involvement of dopaminergic dysfunction in AD pathophysiology.

Our results are consistent with the notion that VTA DA neurons are more vulnerable than their SNc counterparts in preclinical AD models^4,5^, in stark contrast to the opposite association in Parkinson’s disease^77^. While the exact mechanisms underlying preferential pathophysiology are unknown and require further investigation, it may involve differences in the ionic mechanisms that control firing in the two nuclei^78^. In an amyloid-only model of AD, DA neurons from prodromal mice display increased spontaneous and somatically-evoked firing^5^ which may be due to elevated levels of the calcium-binding protein calbindin. Our present data suggest that a decrease in SK channel activity results in hyperactive basal and stimulated DA neuron activity in 3xTg mice. Additionally, our results from WT mice represent the first data exploring intrinsic VTA DA neuron physiology across the lifespan. We previously reported age-dependent decreases in spontaneous and evoked firing along with a delay in rebound firing in the SNc of WT male mice beginning at around 18 months^45,79^. We also reported an apparent decrease in L-type calcium currents, but no change in (presumably SK-mediated) afterhyperpolarization currents with age^45,79^. Consistent with the SNc, here we detected reduced firing frequency (Figure 2a-b) and gain (Figure 2f) in the VTA with age. However, the underlying ionic mechanisms may differ somewhat, as tail currents are decreased in the VTA at 18 months (Figure 3a-c) but do not change with age in the SNc. The difference in age-associated physiological alterations between the two midbrain dopaminergic nuclei suggests that ionic mechanisms are potential factors in subpopulation-specific pathophysiology.

### SK channel dysfunction

SK channel activation regulates both spontaneous firing and burst generation in midbrain DA neurons^14,30,80,81^. During tonic activity, SK channel activation during the ISI sets the timing for the next action potential and dampens sensitivity to synaptic input^41^. SK channel influence on motivated behavior may hinge on their physical proximity to NMDA receptors, where they can serve as a calcium-dependent shunt during excitatory synaptic input^80,81^. SK channels are thought to functionally couple to T-type calcium channels in the SNc^43^, but it is not known whether this holds in the VTA. Roles for SK channels in non-dopaminergic cell types also include synaptic integration^82^ and homeostatic regulation of firing contributing to eyeblink conditioning^83,84^. Taken together, the contributions of SK channels to DA neuron physiology position them as focal points in conditions ranging from drug addiction to neurodegenerative disease^85–88^. Our current work provides a plausible link between SK channel activation and DA neuron physiology in a model of AD.

AD is often characterized as a disorder involving Aβ-dependent synaptic failure^89,90^. Although calcium-activated potassium channels have been implicated in AD-related intrinsic and synaptic pathophysiology^19,91^, until now the specific involvement of CK2 in synaptic alterations within AD has not been described. Notably, CK2 activation reportedly relies on SK channel activity^55^. This implies a potential positive feedback loop, where persistent activity of SK channels caused by increased firing rate or NMDA-dependent activation could decrease the channel’s activity and further potentiate hyperexcitability, which may be consistent with our finding that L-type calcium channel subunit α_2_δ-1 was upregulated in 3xTg DA neurons^92–94^. Furthermore, SK-bound CK2 may serve as a potential link between intrinsic and synaptic imbalances in AD given its proposed role in tuning SK channel calcium sensitivity and thus synaptic integration and homeostatic firing rates. A disparity between firing rate homeostasis and synaptic integration in AD has been proposed as the key mechanism driving the shift from prodromal to clinically apparent stages of AD^95^, and CK2 inhibition abolishes spike adaptation in some neurons^65^, suggesting a novel target for this form of pathophysiological adaptation in AD.

### CK2 inhibition as a potential therapeutic target

SK channels are voltage-independent potassium channels that open in response to a local rise in intracellular calcium. In response to Ca^2+^ binding, CaM undergoes a conformational change that promotes the formation of a complex between CaM and a binding domain on the SK channel^96^. This in turn stabilizes an intrinsically disordered fragment (R396-M412) to open the pore and produce K^+^ efflux^96,97^. This conformational change also depends on PIP2 binding^54^, which is the process promoted by the positive allosteric modulator NS309^54^. Critically, this enhancement is diminished by CaM phosphorylation at T79, the target of CK2. Our Patch-seq experiments identify upregulated CK2 expression in single DA neurons and suggest its involvement in SK channel regulation, as previously reported in other brain regions^65,67^ and *in vitro*^51^. Further, the observed up-regulation of CK2 transcript is consistent with a causative role in the altered firing parameters observed in neurons from 3xTg mice, which we later confirmed using pharmacology. To our knowledge, this is the first report of a role for CK2 in the modulation of DA neuron firing. We also observed a decrease in the effect of NS309 (Figure 4), suggesting a potential hyper-phosphorylation of the SK channel-bound CaM T79 in DA neurons from 3xTg mice.

Direct pharmacological inhibition of CK2 had a dramatic effect in 3xTg DA neurons but did not alter physiological parameters in WT cells (Figure 6). This suggests that VTA DA neurons maintain minimal basal levels of CK2-dependent phosphorylation that then switches to a hyper-phosphorylated state in 3xTg mice. While the exact mechanism triggering increased CK2 activity requires further investigation, CK2 activity is known to be stimulated by Aβ^72^, which accumulates intracellularly in neurons from 3xTg mice^20,98^. We propose that CK2-depedent phosphorylation of SK-bound CaM triggers physiological adaptations in VTA DA neurons of mice that express aberrant levels of Aβ and phosphorylated tau (Figure 7).

**Figure 7.**
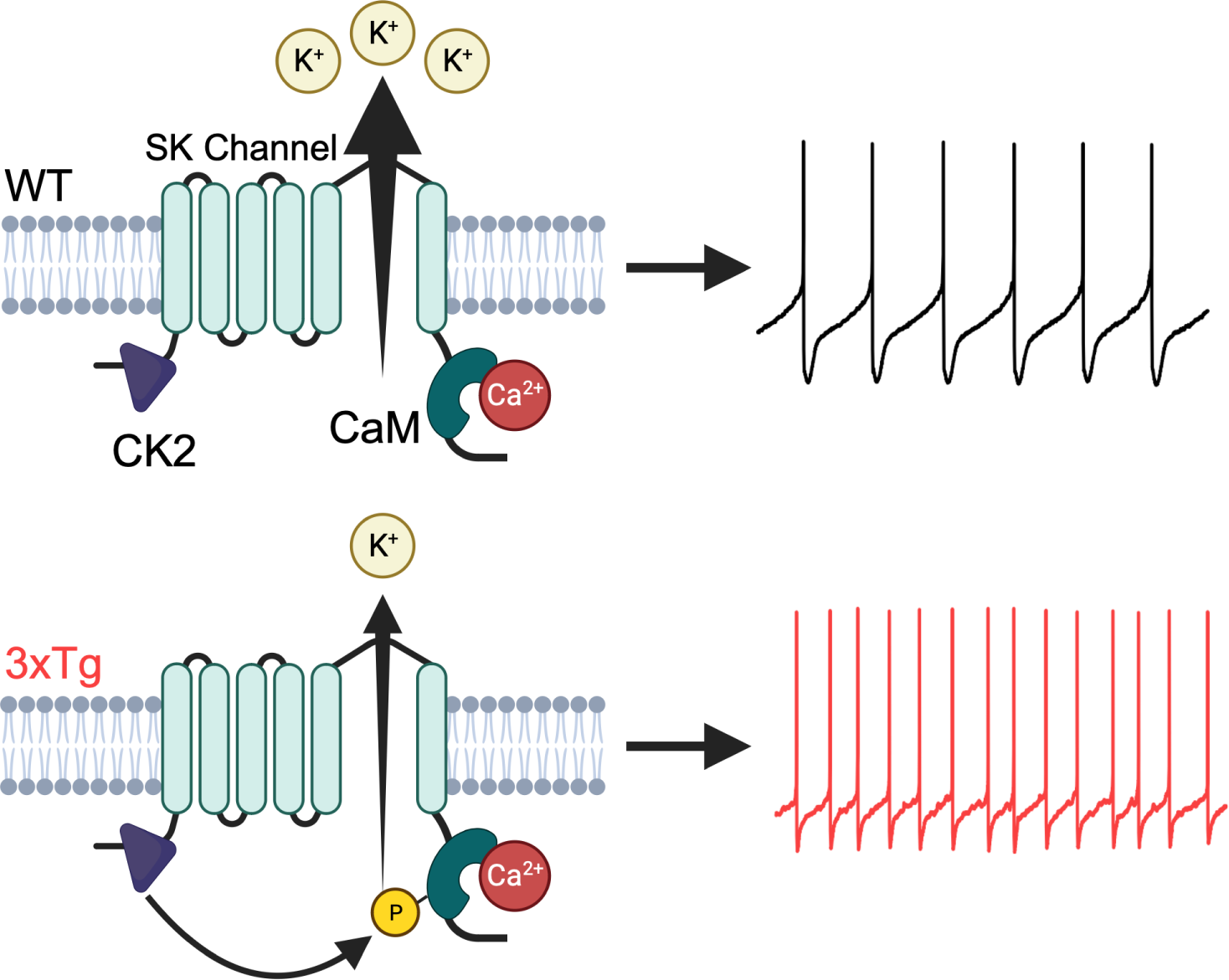
Proposed mechanism of hyperexcitability in 3xTg VTA dopamine neurons. Wildtype mice maintain a balance between CK2 and PP2A resulting in low basal phosphorylation of CaM in DA neurons. In 3xTg mice, hyperactive CK2 results in hyperphosphorylated CaM, effectively decreasing SK channel calcium binding affinity and channel opening. This results in increased firing rate and increased irregularity in 3xTg mice.

CK2 is a pleiotropic kinase. CK2α^-/-^ mice do not develop proper hearts or neural tubes, and mortality occurs mid-gestation^99^. CK2 is also strongly associated with AD^68,71^, PD^100^, and various forms of cancer. The pharmacological profile of CK2 is well described, and clinical trials investigating its antagonist CX4945 are ongoing for basal cell carcinoma, in healthy subjects for tolerability, and for medulloblastoma (ClinicalTrials.gov: NCT03897036; NCT05817708; NCT03904862, respectively). Our data suggest that CK2 is a potential mediator of DA neuron hyperactivity in AD. Pharmacological inhibition of CK2 with SGC rectified irregular firing patterns in 3xTg DA neurons, while application of SGC to WT cells did not result in observable changes in spontaneous firing rate or rhythmicity. This suggests that basal phosphorylation levels may be low in VTA DA neurons, leaving them primed for physiological (or pathological) adaptation. As dopaminergic hyperactivity is a hallmark of multiple conditions, including addiction and psychiatric disorders, these findings point toward a potential role of CK2 in modulating DA neuron firing across diverse contexts.

## METHODS

### Animals

Male and female WT and 3xTg-AD mice were bred in-house at Oklahoma Medical Research Foundation in accordance with the Institutional Animal Care and Use Policies. All mice had *ad libitum* access to food and water. Founders for 3xTg (MMRRC #034830) and WT controls were obtained from Dr. Salvadore Oddo^20^.

3xTg mice were homozygous for all three humanized transgenes (App_Swe_, PSEN1_M146V_, and Tau_P301L_). Due to the hybrid background (C57BL/6;129X1/SvJ;129S1/Sv embryo injected in a B6;129), a separate line on the same hybrid background, not carrying the transgenes, was maintained in parallel and used as control (WT). Breeder genotyping was outsourced (Transnetyx).

### Operant Conditioning Behavior

Food and water were available ad libitum in the home cage at all times for the mice that underwent behavioral testing. Operant testing was conducted in 3-hour daily sessions beginning two hours into the dark cycle in operant chambers that were housed in a sound-dampening cabinet equipped with a fan to minimize external noise. Modular operant chambers (Lafayette Instruments) had two nose-poke holes with a food receptacle between them. For each chamber, one nose poke hole as the correct hole and was illuminated by a stimulus light within it during the session. Responses in the correct hole resulted in reinforcer delivery when the response requirement was met. Responses in the nose poke hole designated as incorrect had no consequence. The correct side was balanced across chambers to decrease side-preference bias. Training began at fixed ratio 5 (FR5); five nose-pokes in the correct hole resulted in administration of a 20 mg palatable food pellet (chocolate-flavored chow pellet, Bio-Serve), activation of a cue-light in the receptacle, and a 5-s tone. There was a 30-s timeout after pellet delivery, where no responding was reinforced, and the stimulus light in the correct side nose poke hole was extinguished. The response requirement was increased across sessions (FR5, FR10, FR20, FR30) when accuracy for the correct side nose poke hole was at least 70% on any single day (accuracy defined as correct side nose pokes/total nose pokes on both sides × 100), with at least 1 day at each response requirement. Acquisition was defined as the first day of training that a mouse obtained 70% accuracy for the correct side nose poke hole. If a mouse did not reach acquisition criterion by the end of the fourteenth session, the mouse was removed from the study and deemed a “non-learner” (Figure 1b). After meeting criterion at FR30 and maintaining stable behavior, mice were subjected to a progressive ratio (PR) session, which was terminated after 60 minutes without earning a reinforcer or 5 hours total. During this PR test session, the FR requirement increased with each pellet earned according to the following schedule: 1, 2, 4, 6, 9, 12, 15, 20, 25, 32, 40, 50, 62, 77, 95, 118, 145, 178, 219, 268, 328, 402, 492, and 603. Breakpoint was defined as the last successfully completed response requirement before the session was ended.

### Brain Slice Preparation

On the day of the experiment, the mouse was anesthetized with isoflurane and decapitated. The brain was removed rapidly and sliced in an ice-cold choline chloride based cutting solution containing the following (in mM): 110 choline chloride, 2.5 KCl, 1.25 Na_2_PO_4_, 0.5 CaCl_2_, 10 MgSO_4_, 25 glucose, 11.6 Na-ascorbate, 3.1 Na-pyruvate, 26 NaHCO_3_, 12 N-acetyl-L-cysteine, and 2 kynurenic acid. Horizontal (200 µm) slices containing the ventral midbrain at the level of the medial terminal nucleus of the accessory optic tract (*mt*) were collected and immediately transferred to a holding chamber containing artificial cerebrospinal fluid (aCSF) containing (in mM) 126 NaCl, 2.5 KCl, 1.2 MgCl_2_, 2.4 CaCl_2_, 1.2 NaH_2_PO_4_, 21.4 NaHCO_3_, and 11.1 glucose. Holding chamber aCSF was supplemented with (in mM) 1 Na ascorbate, 1 Na pyruvate, 6 N-acetyl-L-cysteine, and 0.01 MK-801, to minimize excitotoxicity associated with slicing. Slices recovered for thirty minutes at 32°C followed by an additional 30 minutes or more at room temperature prior to recording. For the SGC-preincubation experiments, 1 µM was added to the holding chamber, and slices were maintained in this solution for 1-3 hours prior to recording. Neurons were recorded up to ten hours after slicing.

### Electrophysiology

Slices were transferred to a recording chamber where they were perfused with warmed aCSF (Warner Instruments inline heater, 32-34°C) at a rate of 2 mL/min via gravity or a peristaltic pump (Warner Instruments). Slices were visualized under Dödt gradient contrast (DGC) optics via an upright microscope (Nikon). Putative VTA dopaminergic neurons were identified first by location (>100 µm medial to *mt* at the level of the fasciculus retroflexus [*fr*]). A subset of recorded neurons were labeled with biocytin and imaged on a confocal microscope (Zeiss LSM 710) or imaged under DGC on the rig for localization (Extended Figure 4). Electrophysiological criteria for VTA dopamine neuron identification included slow, spontaneous firing in the cell-attached configuration (0.5-8 Hz) and wide (>1.1 ms) spikes^101^, inactivation time constants of the A-type potassium current between 30 and 300 ms^36^, and a long (>250 ms) rebound delay after a hyperpolarizing step (−100 pA) in current clamp^34,35^. SNc dopamine neurons were recorded <50 µm from *mt* (Extended Figure 1a), and electrophysiological identification parameters included slow (1-5 Hz) spontaneous firing and wide (>1.1 ms) spikes in the cell attached configuration, fast-inactivating A-type potassium conductance, and hyperpolarization-activated inward current (I_H_) > 100 pA. For cell-attached and whole-cell recordings, patch pipettes were pulled from thin-wall glass (World Precision Instruments or Warner Instruments) and had resistances of 2.5-3 MΩ when filled with an intracellular solution containing (in mM): 135 K gluconate, 10 HEPES, 5 KCl, 5 MgCl_2_, 0.1 EGTA, 0.075 CaCl_2_, 2 ATP, 0.4 GTP, pH 7.35^102^. Whole-cell internal solution had a liquid junction potential of −17.1 mV, which was calculated with the stationary Nernst–Planck equation using LJPcalc software (https://swharden.com/LJPcalc) and corrected offline. Perforated-patch pipettes were pulled from standard wall glass (Warner Instruments) and had resistances of 3.5-4.5 MΩ. The tip was first filled with (in mM) 140.5 KCl, 7.5 NaCl, 10 HEPES (pH 7.0) and then backfilled with the same solution containing 1-2 µg/mL gramicidin-D. Recordings generally began when perforation was adequate to display action potential overshoot of 0 mV, and recordings were terminated if break-in occurred. Patch-seq pipettes were pulled from thin-walled glass similar to whole-cell recording pipettes except with a slightly smaller tip (3-4 MΩ) to allow efficient nucleated patch formation^57^. All pipettes were pulled on a horizontal PC-10 or PC-100 puller (Narishige International).

### Chemicals and Pharmacological Agents

NS309 was purchased from Alomone Labs. Apamin was purchased from Echelon Biosciences. SGC-CK-2 was purchased from Tocris. All additional chemicals were purchased from Sigma-Aldrich.

### Electrophysiological Data Acquisition and Analysis

Recordings were acquired with a MultiClamp 700B amplifier and digitized with an Instrutech ITC-18 board. AxoGraph (v1.7.6 and v1.8.0) and LabChart (ADInstruments) software were used for recording. Voltage-clamp recordings were low-pass filtered online at 6 kHz and digitized at 20 kHz. Current clamp recordings were low-pass filtered at 10 kHz and digitized at 20 kHz. All analyses were conducted offline via custom scripts in Python (3.9). Code used for analyses will be made publicly available following publication (https://github.com/heblanke/DA_neuron_analysis). Cell-attached recordings were filtered using a Savitzky-Golay filter and action currents were detected in the negative direction. In current clamp, spikes were generally detected at the upward crossing of −10 mV, with the criterion that dV/dt must exceed 10 mV/ms. ISI times and voltages were collected by subtracting the first spike time from the times between each pair of spikes and were resampled at 1000 points spanning each ISI to allow for averaging across ISIs and across different cells. The minimum voltage value was taken as the minimum of the average ISI for a cell. The average voltage across the mAHP was calculated as the average voltage from 10 ms after the minimum point to 100 ms after the minimum point of the ISI. The ISI voltage trajectories illustrated are grand averages across all cells in a group. Tail current AUC was calculated using the trapezoidal rule implemented in NumPy. A-current inactivation time constants were generated by fitting a bi-exponential curve to currents using the SciPy curve fit function. Apparent threshold values were determined by averaging spike voltages for a given cell and determining the voltage at which 10 mV/ms was crossed. Action potentials were binned into 10 ms windows to measure frequency and CV of ISI in order to avoid confounds of rate drift. Representative traces of action potentials were truncated at either 10 mV (Figures 2, 3, and 6) or at 20 mV (Figure 4).

### Patch-seq

Patch-seq required slight modifications to recording parameters to minimize confounds in both electrophysiological recordings and sequencing. Patch-seq internal was a modification of whole-cell internal solution containing (in mM) 124 K gluconate, 10 HEPES, 5 KCl, 5 MgCl_2_, 0.1 EGTA, 0.075 CaCl_2_, 20 µg/mL glycogen, and 0.5 units/µL RNase inhibitor (M0314L, New England Biolabs). The voltage step protocol used to measure the tail current was unchanged. To minimize recording time before cell extraction, additional electrophysiological parameters were not recorded. After recording was complete, light negative pressure was applied to the pipette until the nucleus was visible at the tip of the pipette, or for approximately three minutes. Next, the recording pipette was slowly retracted along the Z-axis. Successful extractions included an obvious nucleated patch^57^. After the nucleus was visualized above the slice, the pipette was rapidly retracted from the bath, and the cellular contents within the pipette were deposited in a 0.2 mL PCR tube containing 0.5 µL of NEB Next lysis buffer and 0.25 µL NEB murine RNase inhibitor. Tube contents were then rapidly spun down and flash frozen on dry ice. Samples were stored at −80°C until RNAseq libraries were generated.

### RNASeq Library Generation

Directional libraries were constructed using NEBNext Single Cell/Low Input RNA Library Prep Kits from Illumina following manufacturer instructions. In brief, poly-adenylated mRNA was magnetically captured and then eluted from the oligo-dT beads. mRNA was then fragmented and reverse-transcribed, and the cDNA was purified using SPRISelect beads (#B23318, Beckman Coulter). Purified cDNA was eluted in 50 μL 0.1 × TE buffer. Adaptors were diluted and ligated to cDNA, and the cDNA was amplified by 28 cycles of PCR. SPRISelect beads were used to select for proper library size. Purified library quality was then assessed on a tapestation (#4200, Agilent Technologies) using HSD1000 screentapes (#5067–5584; Agilent Technologies). Due to the high number of amplification cycles, occasionally libraries required additional rounds of SPRISelect purification. Qubit 1 × dsDNA HS Assay kit (#Q33230, Thermo Fisher Scientific) was used to quantify libraries. Libraries were pooled and sequenced 2x 100 bases on an Illumina MiSeq. Fastq data will be made available on the gene expression omnibus (GEO) upon acceptance.

### RNASeq Analysis

Reads were trimmed and aligned before differential expression statistics and correlation analyses in Strand NGS software package (v4.0; Strand Life Sciences). Samples with reads were aligned against the full mm10 transcriptome RefSeq build (2013.04.01). Alignment and filtering criteria included the following: adapter trimming, fixed 5-bp trim from 5′ and 3′ ends, a maximum number of one novel splice allowed per read, a minimum of 90% identity with the reference sequence, a maximum 5% gap, and trimming of 3′ ends with Q < 30. All duplicate reads were then removed. Normalization was performed with the DESeq2 algorithm. Transcripts with an average read count value ≥ 1 in at least 30% of the cells in at least one group were considered expressed at a level sufficient for quantitation. For statistical analysis of differential expression, a t-test with Benjamini–Hochberg multiple testing correction (BHMTC) was performed with a False Discovery Rate cutoff of <0.1. For those transcripts meeting this statistical criterion, an absolute fold change > 1.25 cutoff was used to eliminate those genes that were statistically significant but unlikely to be biologically significant and orthogonally confirmable because of their very small magnitude of change. Visualizations and principal component analyses (PCAs) were performed in Strand NGS (version 4.0) and WebGestalt 2019^103^. Gene set enrichment analysis (GSEA) was performed using GSEA v4.3.2.

### Statistical analysis

Statistics on electrophysiological data were analyzed in Prism (versions 9 and 10), and statistics and calculations for RNAseq were conducted in Python and Prism. All electrophysiology parameters collected were batch analyzed to minimize bias. Unpaired and paired two-tailed t-tests were used for two-group comparisons. Two-way ANOVAs with Sidak’s post-hoc test for multiple comparisons were conducted where indicated. All error bars represent standard error of the mean (s.e.m.) unless otherwise noted.

## Acknowledgements

This work was supported by grants from the NIH (R21 AG072811 and R01 AG052606 to MJB, F31 AG079620 to HEB, P30AG050911 and R01AG059430 to WMF), the Department of Veterans Affairs (I01 BX005396 and IK6BX006033), the Presbyterian Health Foundation, and the Oklahoma Center for Adult Stem Cell Research, a program of TSET. We would like to thank Dr. Ezequiel Marron for insightful questions about the project, and Drs. Stephanie Gantz, Christopher Ford, and Matthew Higgs for their invaluable comments and suggestions.

