## Extended Figures for "VTA dopamine neurons are hyperexcitable in 3xTg-AD mice due to casein kinase 2-dependent SK channel dysfunction"

**Extended Figure 1. SNc DA neurons are unaffected in 3xTg mice. a.** Schematic representing a subset of recoded neurons, indicating location of recorded cells in the SNc. **b.** Representative five-second traces of cell-attached recordings in WT (black) and 3xTg (red) SNc DA neurons. No difference in firing rate (**c.**) or CV of ISI (**d.**) between the two genotypes was detected. **e.** Traces representing evoked firing at +50 pA in the whole-cell configuration for WT (black) and 3xTg (red). **f.** Averaged frequency-current curves for WT and 3xTg indicate no change in evoked firing. **g.** No difference in the gain (slope of the steady state [0-60 pA] F/I curves) between WT and 3xTg neurons was observed. **h, i.** Tail current charge was unaffected in 3xTg SNc neurons. **j, k.** A-type potassium current peak amplitude was also unaffected.

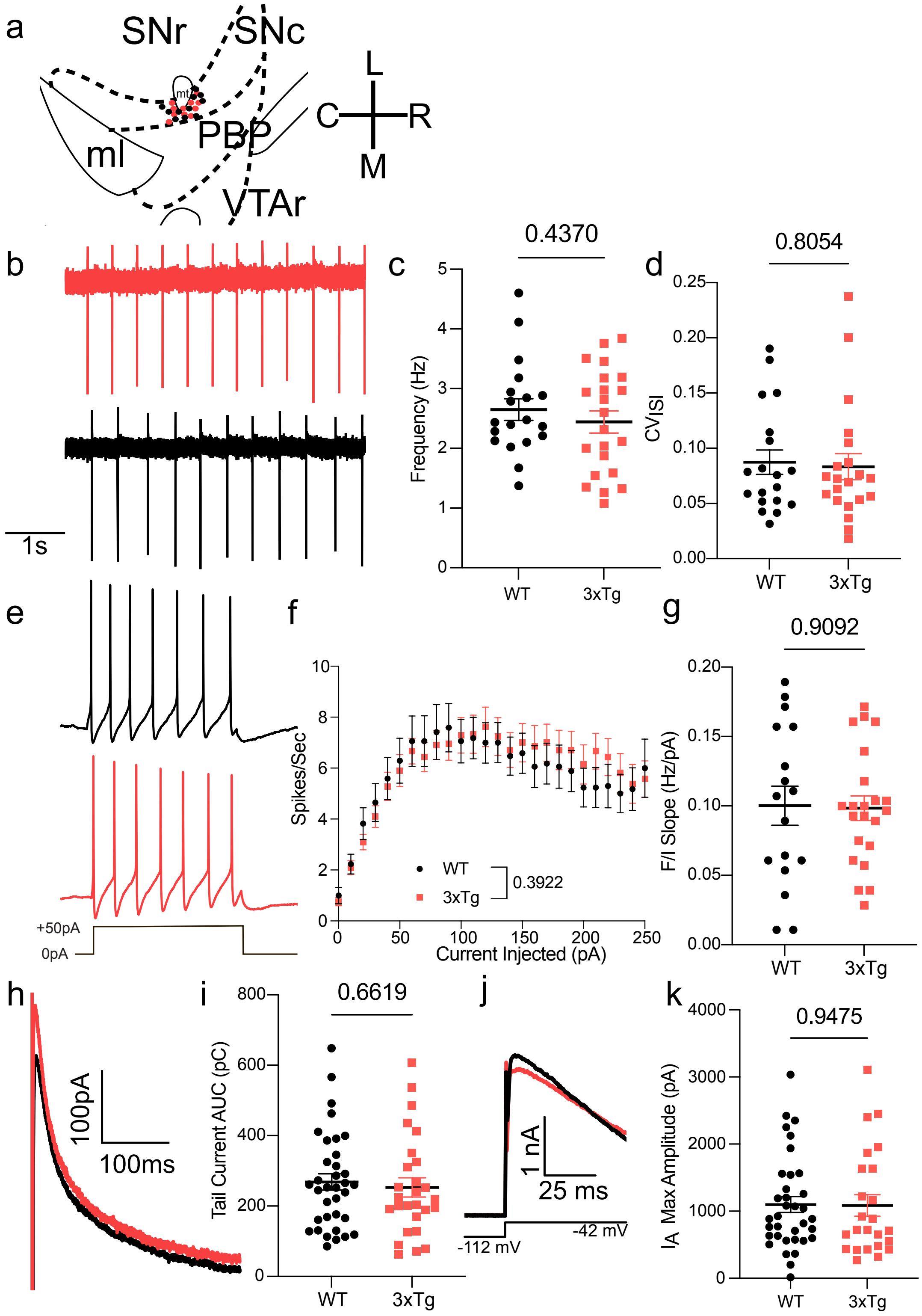

**Extended Figure 2. The tail current in VTA DA neurons is mostly apamin-sensitive. a**. Representative traces before (black) and after (orange) apamin (100 nM). **b**. Tail current maximal amplitudes, evoked by a step from (-72 mV to -17 mV). Apamin was washed on for ~18 minutes and did not wash out within the duration of these recordings. **c**. Tail current maximal amplitudes before (average of the first 7 sweeps) and after (average of the final 7 sweeps) apamin (paired two-tailed t-test, P=0.0015).

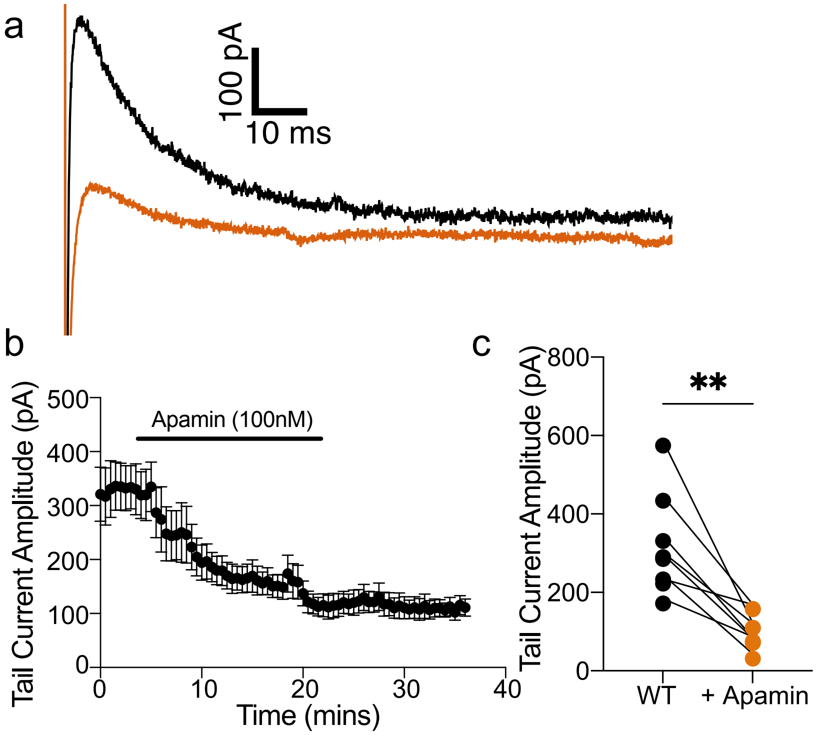


**Extended Figure 3. 12 mo 3xTg DA neurons exhibit slow rebound delays but unaltered A-type potassium currents. a**. Representative trace of A-type current in 12 mo WT (black) and 3xTg (red) neurons (step from -72 to -112 to -52 mV)[40]. **b**. $I_A$ maximal amplitude is not different by genotype but does display an age x genotype interaction (N = 296, $F_{genotype\ x\ age}$ = 4.319, P=0.0053). **c**. The inactivation time constant is also unaffected by genotype but displays an age x genotype interaction ($F_{age}$ = 3.083, P=0.0277; $F_{age\ x\ genotype}$ = 3.039, P=0.0294).

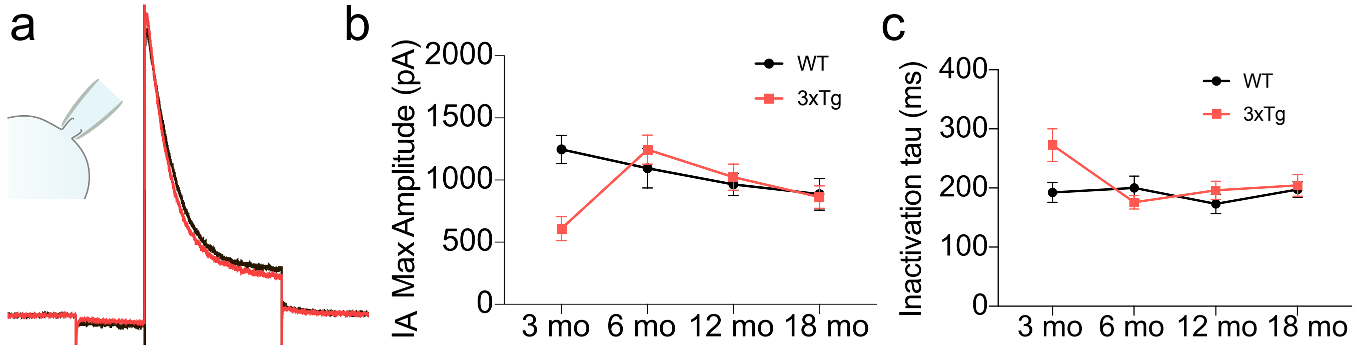

**Extended Figure 4. Recorded VTA dopamine neurons were predominately located in the parabrachial pigmented nucleus (PBP).** A subset of recorded cells was labeled with biocytin in the recording pipette and imaged on a laser scanning confocal microscope. For some cells, location relevant to the medial terminal nucleus of the accessory optic tract (*mt*), the medial lemniscus (*ml*), and fasiculus retroflexus (*fr*) was noted while recording to allow for approximate mapping to the Paxinos and Franklin 2019 stereotaxic mouse brain atlas. All cells were located ≥100 µm medial to *mt,* rostral to *ml,* and lateral to *fr.* DA neurons in this part of the VTA are heterogeneous, and we aimed to capture a representative sample of the subtypes present in this region[4].

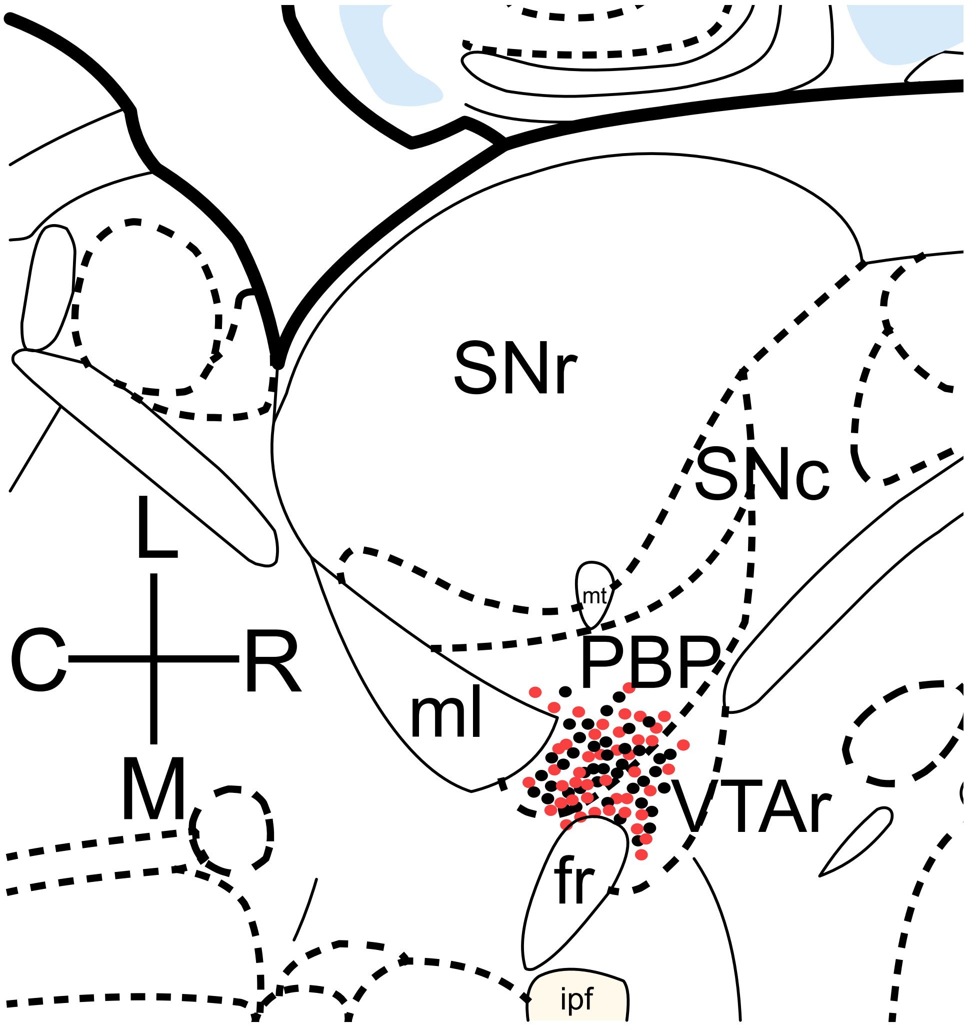
